# Ancient DNA supports that the evolution of Skin Tone in East Asians intensified after their split with Native Americans

**DOI:** 10.1101/2023.09.03.553077

**Authors:** Manuel Ferrando-Bernal

## Abstract

Skin tone has been deeply studied in European populations both using modern and ancient DNA. However, other populations are underrepresented in such studies. One such population is East Asians, for which, interestingly, it has been claimed to evolve light skin tones in parallel to Europeans. Moreover, it is not clear whether this happened before or after their split from Native Americans. Over the last few years, several studies have sequenced hundreds of ancient genomes belonging to East Asians ancient populations. Additionally, some variants have been associated with light skin in East Asians. To shed some light, I applied a Polygenic Risk Score for some of the variants associated with light skin, in 237 modern Native Americans and East Asian individuals and in more than 700 East Asians ancient samples. The results suggest that this phenotype may have started to evolve in the ancestors of East Asians and Native Americans but intensified after their split.

## 1. Introduction

Differences in skin tones are primarily attributed to a complex interaction of several genes and environmental factors, such as ultraviolet (UV) radiation exposure (1). This polygenic phenotype kept the attention of evolutionary biologists and anthropologists for decades as there is a gradient in the colour among human populations that correlates with latitude (2; 3).

The predominant factor influencing skin tone is the production of melanin, a pigment that provides coloration to the skin, hair, and eyes. There are two main types of melanin: eumelanin, which is responsible for darker pigmentation, and pheomelanin, which contributes to lighter pigmentation. East Asians and Europeans populations generally have lower levels of eumelanin and higher levels of pheomelanin compared to populations with darker skin tones. The specific genetic variations associated with these differences in melanin production have been studied extensively, and several genes have been identified as having an impact on skin pigmentation. But, most of these studies have been carried out with individuals of European ancestry, recovering more than 170 single nucleotides variants associated with this phenotype (4). Contrary, only a few variants associated with light skin have been identified in East Asian populations.

Ancient DNA studies have revolutionised the way we can understand the past, confirming questions such as if we interbreed with Neanderthals and Denisovans (5; 6), and the tempo for the evolution of some specific traits such as lactose tolerance capacity in some Europeans (7). Some of these studies suggest that selection for light skin in Europeans intensified during the Neolithic, but may have started thousands of years before (1). Light skin seems to have evolved convergently in East Asians (8; 9), and it has been suggested that it happened before their split with Native Americans (10), which may have happened probably around 20,000 ya. Contrary, it has also suggested that it may have evolved after the first populations moved towards the American continent (11). Furthermore, models using modern-day samples date it to between 35,000 and 5,000 ya, likely around the las 10,000 ya (10) and unfortunately, no one has studied it using ancient samples to date. Here, I tried to give more concise dates for the evolution of this phenotype. To do so, I recovered several Single Nucleotide Polymorphisms (SNPs) related to light skin in East Asians and applied a Polygenic Risk Score (PGR) for light skin analysis in 200 modern and 745 ancient East Asians encompassing the last 40,000 ya. These results show that depigmentation skin may have evolved during the last 15,000 ya., sometime later than the split among East Asians and Native Americans.

## 2. Material and methods

### Databases

#### SNPs related to light skin

Most GWAS have focused on the European populations, allowing the recovery of more than 170 SNPs related to light skin (1;4). However, only few studies have been applied with East Asians individuals. In this study, I use the most recent article with 6 SNPs related to light skin in East Asians (12) and in which they also give the beta values for the GWAS association. I additionally add the SNP rs1800414 that has been identified as related to light skin evolution in East Asians multiple times (8; 9). As this SNP is not among the 6 SNPs recovered by (12), I need to correct its betta value. To do so I use one the 6 SNPs, rs74653330, which were associated together with rs1800414 in (13) (See Supplementary Table 1).

#### Availability of Data and imputation

All modern and ancient genomes (1240k) were downloaded from https://reich.hms.harvard.edu/allen-ancient-dna-resource-aadr-downloadable-genotypes-present-day-and-ancient-dna-data. They were then filtered for East-Asian samples (belonging to South Korea, Japan, Mongolia, China or Taiwan) or Native Americans (belonging to Karitiana, Piapoco, Pima and Surui populations, as that have been shown to not have admixed ancestry from European or African origin). A total of 1003 individuals were downloaded (31 modern Native Americans, 206 modern East Asians and 766 ancient East Asians). The data was turned from the eigenstrat format into plink format and to VCF format. To do so I used the R library ADMIXTOOLS to transform from the Eigenstrat format to plink with the function “eigenstrat_to_plink”. Then I used PLINK 1.9 to transform the data into VCF format with the function “recode vcf”. The resulting VCF file was standardised with bcftools (version 1.14) by matching chromosome notations and REF/ALT allele order to 1KG. Chromosomes were renamed with the bcftools function --rename-chrs. Alleles were matched with bcftools norm “--check-ref ws --fasta-ref <GRCh37_ref_fasta> --do-not-normalize” and then split the file into chromosome VCFs. Conform-gt (version 24May16.cee) was used to make the VCF files consistent with 1KG reference. For this, the reference genotype data was dowloaded from https://bochet.gcc.biostat.washington.edu/beagle/1000_Genomes_phase3_v5a/b37.vcf/. 84420 SNPs (∼7%) in the East-Asians VCFs did not match with the 1KG reference and were removed from output VCFs. The resulting VCF files were phased and imputed with Beagle 5.4 (version 22Jul22.46e). To phased them I used 1000 genome phase 3, downloaded as reference panels from https://bochet.gcc.biostat.washington.edu/beagle/1000_Genomes_phase3_v5a/b37.bref3/ and GRCh37 genetic maps from https://bochet.gcc.biostat.washington.edu/beagle/genetic_maps/

#### Polygenic Risk Score estimation

Polygenic Risk Scores are applied to study phenotypes that are influenced by multiple genomic variants (14-18). In humans, examples of these phenotypes are skin pigmentation, bone mineral density or height. A genetic risk score can be obtained by summing all the alleles related to a particular phenotype (15) associating a value to the likelihood to develop a particular phenotype (for example, developing a higher or lower bone mineral density, or to develop a darker or a pale pigmentation skin). Polygenic risk scores have been applied several times in ancient DNA studies, for example, to study the height in ancient Europeans (16, 17) to study the genomic variants associated with a bone disease (14), to study the evolution of attention/deficit hyperactivity disorder in Neanderthals and ancient Europeans (18) and, as here, the likelihood to develop a particular skin pigmentation, but in Europeans (1). In this study, the genetic risk score was calculated by summing in each individual all the alleles related to skin pigmentation multiplied by their betta value (a value that measures the intensity that the alleles are related to skin pigmentation) <u>See Supplementary Table 1</u>).

#### Statistical analysis

Principal component analysis (PCA) is one of the most commonly used statistical tools. Is it used to analyse multivariate data so it has been broadly applied to genomics datasets, including dozens of aDNA studies. PCA transforms a number of correlated variables into a smaller number. Large genomic datasets, such as the one used here, are composed of hundreds of thousands of SNPs and each SNP gives information on how the individuals of the dataset can be distributed. The PCA analysis summarises this information by accounting for correlations among these SNPs. For example, populations that share a particular or a recent ancestry will have hundreds or thousands of SNPs correlating together, and all this information can be pulled together into a single variable, thus reducing the number of variables able to explain the information in the database and facilitating tools to visualise it. For example, it is common to observe in a PCA plot, the PC1 in the x-axis and the PC2 in the y-axis, giving an idea of how the individuals in the database cluster together. As related individuals will share more alleles among them, they will cluster together in a PCA analysis. So, here I applied this methodology to calculate relationships among the modern and ancient individuals in the database. The PCA analyses were calculated in PLINK (19) and then plotted in R Studio with ggplot2 (21).

The Kruskal-Wallis test is a non-parametric method commonly used to test for differences among populations. It is used to compare two or more dataset with equal or different sizes (as is the case here). I used the option in R studio to see differences in the PGR among the 19 modern East Asians populations.

Pearson correlation coefficient is a method used to calculate linear regression among two datasets. Conversely, Spearman correlation is used to find correlations that can but do not need to be linear. Here I applied both to test for correlations among the PGR of the modern East Asians and environmental variables from where each individual lives. Both tests were applied using their option in R studio.

#### Environmental Data

The coordinates of the individuals were obtained from the David Reich lab. database (https://reich.hms.harvard.edu/allen-ancient-dna-resource-aadr-downloadable-genotypes-present-day-and-ancient-dna-data). Solar radiation was obtained from the Global Solar Atlas webtool (https://globalsolaratlas.info/support/about). Annual, winter and summer UV-radiation levels were downloaded from the NASA Prediction of Worldwide Energy Resource (POWER)(https://power.larc.nasa.gov/data-access-viewer/).

## 3. Results and discussion

### 3.1 Differentiation among populations in their Polygenic Risk Score for skin tone cannot be explained by differences in their ancestry

To test differences among modern populations I conducted a boxplot with the Polygenic Risk Score for skin tone among 19 modern East Asians populations (Figure 1A, see Supplementary table 2). I found that populations statistically differ among them in the PGR (Kruskal Wallis test, p value 0.001899). Specifically, Oroqen, Xibo, Atayal and the Korean population show the highest mean values in the PGR, pointing toward a possibly slightly lighter pigmentation (despite them showing an important variation in their skin tone inside each population). On the other hand, the Lahu population presents the lower value of PGR, suggesting a possible toned skin pigmentation. I consider that it is important to highlight that despite I am referring to the populations with high PGR as possible dark skin, these population may have evolved their particular light skin alleles, as for example the Uyghur who seem to have inherited pigmentation alleles from Indo-European populations (22) and which is the population with more diversity in their PGR. Also, it is important to highlight that most of the SNPs used here were obtained in a GWAS from Korean populations, which may explain why Koreans seem to be one of the populations with the highest PGR score.

**Figure 1.**
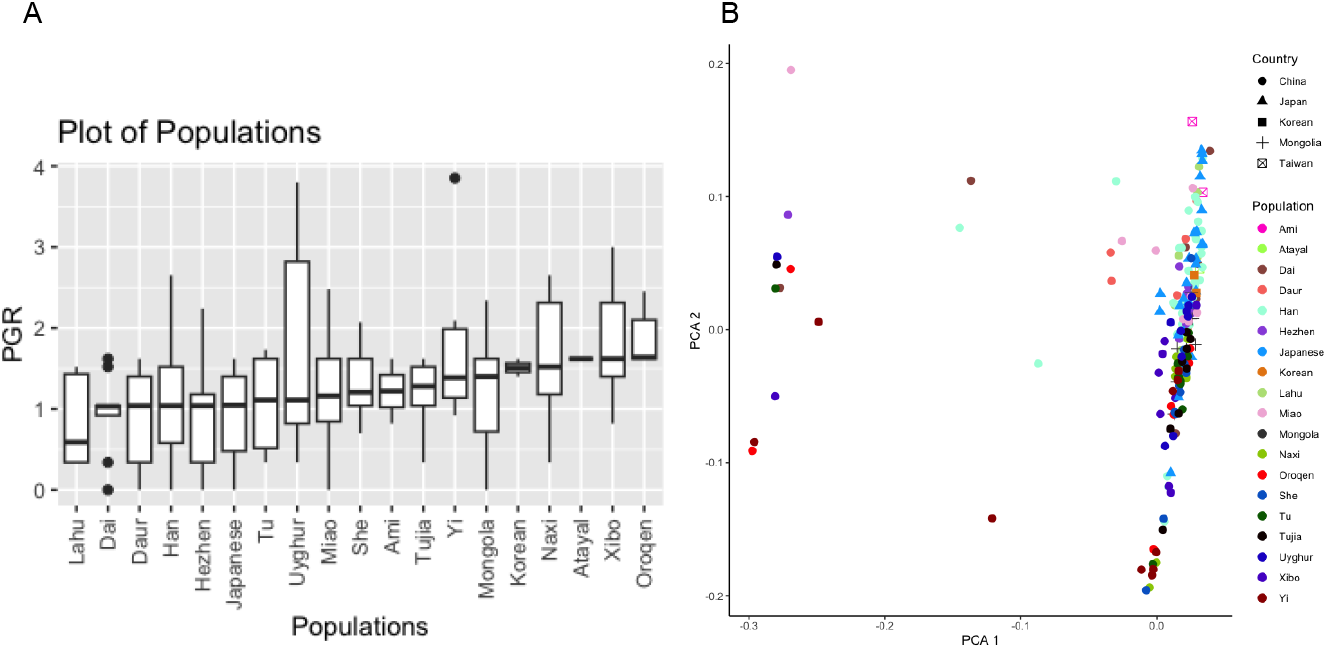
(A) Boxplot of the PGR values in 19 modern East Asians populations. The x-axis represents the Polygenic Risk Score (PRS), and the y-axis represents each population. (B) PCA with 206 modern East Asians individuals. The x-axis represents the Principal Component 1, the y-axis represents the Principal Component 2. Each dot represents an individual. Different shapes represent the country where the individual lives. Most of the individuals cluster together whereas some individuals cluster separately to the main cluster. These individuals are: three Han, three Yi, two Miao, two Oroquen, two Daur, two Dai, one Hezhen, one Uyghur, one Mongol, one Tu and one Xibo individuals. No Lahu individual clusters separately to most of the individuals in the dataset.

It is possible that we are observing an adaptation to differences in the environmental cues (i.e. sun radiation at which each population is exposed or geographical variables such as latitude or longitude). However, there is also the possibility that differences in the allele frequency among individuals are caused by differences in their ancestry (1, 14). For example, if individuals with higher values for a less pigmented skin genetic score belong to a different population than individuals with lower values they will be sharing different alleles, including those associated with skin tone. Specifically, I decided to test it for the Lahu individuals as it is the population with the lowest PGR values, and the Oroquen, Xibo, Atayal and Korean individuals as are the populations with the highest mean PGR values. Interestingly, Lahu (and Dai which is the second population with the lowest mean PGR value) is an ethnic group that shows close genomic similarities with other southerners such as Cambodian and Vietnamese populations (22). This is due to Lahu inhabiting mainland SouthEast Asia, where native populations tend to have a darker pigmentation in the skin tone. All this opens the possibility for a possible gene flow among the Lahu inhabiting China and the Lahu inhabiting Thailand and Vietnam, which may explain the differences in their PGR. To test this, I conducted again a Principal Component Analysis (see Figure 1 B; see “Material and methods”) with all the modern East Asian individuals. I observed that most of the individuals cluster together, with some individuals cluster separately. These individuals are: three Han, three Yi, two Miao, two Oroquen, two Daur, two Dai, one Hezhen, one Uyghur, one Mongol, one Tu and one Xibo individuals. Interestingly, no Lahu individual clusters separately to the main cluster, thus suggesting that differences in their ancestry cannot explain differences in the PGR values in the Lahu individuals. Regarding the populations with highest PGR values only two Oroquen (from 9 individuals) and one Xibo (from 9 individuals) cluster separately to the main cluster, thus suggesting that their highest values cannot be explained only by differences in their ancestry.

### 3.2. Correlation of Polygenic risk score for skin pigmentation with geographical variables

It is commonly accepted that light skin evolved in Europeans as an adaptation. UV radiation is needed to synthesise vitamin D, a vitamin needed for a correct skeletal and immune system development. Melanin blocks most of the UV radiation and, as this radiation decreases as the latitude increases, light skin pigmentation may have evolved to allow the synthesis of vitamin D in the high European latitudes. The same hypothesis has been suggested for East Asians, but with low evidence to date. To test for it, I study for a correlation among PGR the geographical location in the modern East asians populations. These results found that PGR highly correlates with longitude (Pearson correlation -0.24, p value 0.00036) but not with latitude (See Figure 3). Interestingly, sun radiation highly differs in Eastern Eurasia correlating higher radiations with lower longitudes and with high latitudes (23).

Additionally, I test for correlations among PGR and the sun radiation at the location where each modern individual lives. I do not find statistical correlation among annual mean UV radiation and the PGR, nor with the radiation levels during the winter months (December and January). However, the correlation increases only with the radiation UV levels during summer months (June, July and August), becoming statistically significant specifically with the UV-A radiation (Pearson correlation 0.149, p value 0.031; see Table 1; <u>Supplementary table 3</u>).

**Table 1.** Table summarising the correlation among PGR and different Sun radiation features.

|  | **Correlation** | **Pearson p value** | **Correlation** | **Spearman p value** |
| --- | --- | --- | --- | --- |
| **Solar radiation** | 0.09261448 | 0.1855 | 0.108137 | 0.1218 |
| **Mean Annual UV-radiation** | 0.05741339 | 0.4124 | 0.03462143 | 0.6213 |
| **Mean Annual UV-radiation** | 0.01613903 | 0.8179 | 0.04289939 | 0.5404 |
| **UV-A radiation during summer** | **<u>0.149894</u>** | **<u>0.03152\*</u>** | 0.106871 | 0.1263 |
| **UV-B radiation during summer** | 0.1240274 | 0.07571 | 0.06645975 | 0.3426 |
| **UV-A radiation during winter** | -0.04610212 | 0.5105 | -0.005982569 | 0.932 |
| **UV-B radiation during winter** | -0.02005814 | 0.7748 | -0.03988811 | 0.5692 |
| **Latitude** | 0.07768953 | 0.267 | 0.04664504 | 0.5056 |
| **Longitude** | **<u>-0.2464407</u>** | **<u>0.0003559\*</u>** | **<u>-0.1260844</u>** | **<u>0.07094</u>** |

### 3.3. East Asians light skin likely started to evolved after the split with Native Americans

One of the most intriguing questions related to the evolution of less pigmented skin in East Asians is if it happened after or before their split with Native Americans (10). The ancestors of Native Americans entered via the Alaska peninsula into North America after their split with ancient East Asians, not before 15,700 ya, likely around 20,000 ya (24, 25). If depigmentation skin evolved in the ancestors of those two populations, Native Americans should show similar PGR for skin pigmentation than East Asians. To test it, I calculated their PGR for skin tone and compared them. I found that the distribution of the PGR in the Native American populations falls inside the distribution of most of the East Asians populations (see Supplementary Table 5, Figure 2), but with a PGR suggesting a higher tone than most of the East Asians populations (except from the Lahu). Despite these results suggesting that depigmentation of the skin tone in East Asians may have evolved after the split among Native Americans and East Asians, I decided to conduct additional analysis.

**Figure 2.**
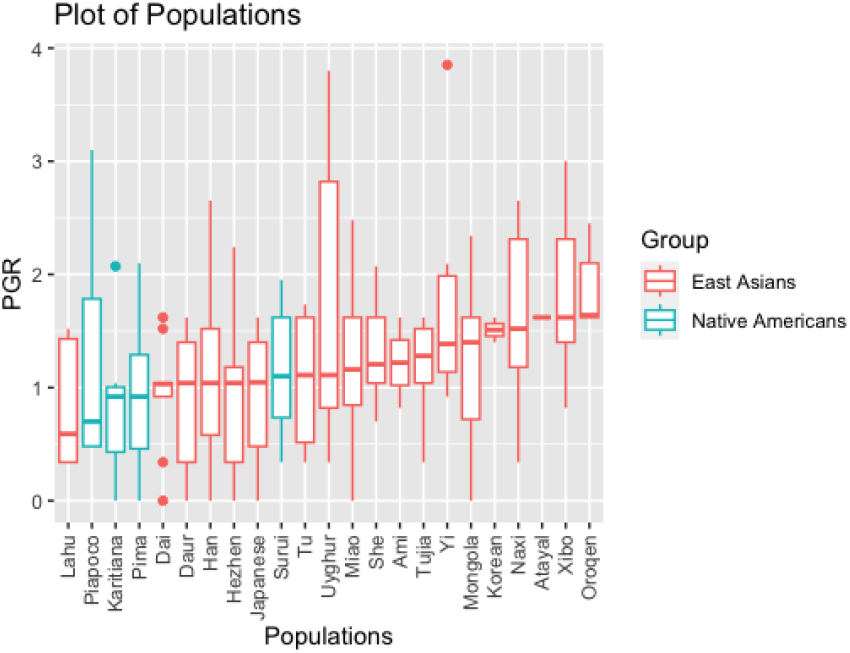
Boxplot with the PGR distribution among the modern East Asians and Native American populations. In the x axis are plotted the PGR values and in the y axis are plotted the populations. Native American populations are highlighted in green and the East Asians populations are highlighted in orange. The mean PGR values of the Native American populations are among the lowest of most of the East Asians.

To confirm it, I test it by studying the temporal changes in the PGR during the last 40,000 ya. As I am exploring the evolution of depigmentation skin in East Asians, I need to generate a database of modern and ancient individuals that may be related. To do so I conducted again a Principal Component Analysis, but this time with the 972 modern and ancient East Asians individuals (see Figure 3; see “Material and methods”, see <u>Supplementary Table 4</u>) .

**Figure 3.**
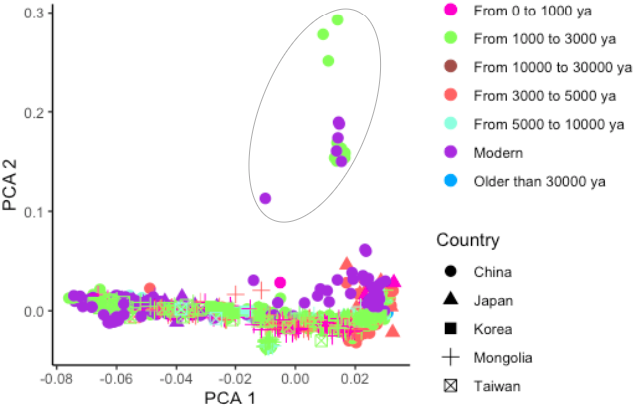
PCA with 972 ancient and modern East Asians individuals. Each dot is an individual. Different shapes represent the country where the samples were recovered and each colour represents the time period where the individuals lived. Most of the individuals cluster together (including several modern and ancient) whereas a few of them cluster separately (highlighted by surrounding with a circle). For further analysis I decided to exclude these individuals that cluster separately. 945 individuals are reminded for further analysis.

In the PCA we can observe two main differentiated clusters. One represents most of the individuals in the dataset, including some modern and all ancient samples from Japan, South Korea, Taiwan and Mongolia. Some modern individuals from China (including three Naxi individuals from the Himalaya and the three Oroquen people, inhabitants of Northern China and inner Mongolia) as well as some individuals that lived among 1,000 and 3,000 y.a. appear to cluster separately. For the further analysis, I decided to discard those individuals. 200 modern and 745 ancient individuals remain for further analysis.

These 945 East Asians encompass a transect of 40,000 years of inhabitation in East Asia. Most of the individuals need to be imputed previously to the PGR estimation (see Materials & Methods). Once I have their PGR I plot it against the individual age to see the evolution of this phenotype through time. The results suggest that light skin may have evolved after the split among East Asians and Native Americans (see Figure 4A, see Supplementary Table 6). For example, all ancient individuals older than 15,000 ya show low values in their PGR, pointing towards they carried alleles not related with light skin in modern East Asians. These individuals include one dating from 19,427 ya (around the age when Native Americans and East Asians split (24, 25)) probably had a darker skin tone than modern East Asian individuals. This individual lived in what is now called Heilongjiang, at the extreme North East of China, close to the Bering Strait from which ancestors of Native Americans crossed. Despite other studies indicating that the ancestors of Native Americans and East Asians possibly had some light skin tone (10), the results here suggest that most of the related light pigmentation skin alleles increased their frequency in the last 15,000 ya (see Figure 4A, see <u>Supplementary Table 5</u>).

**Figure 4.**
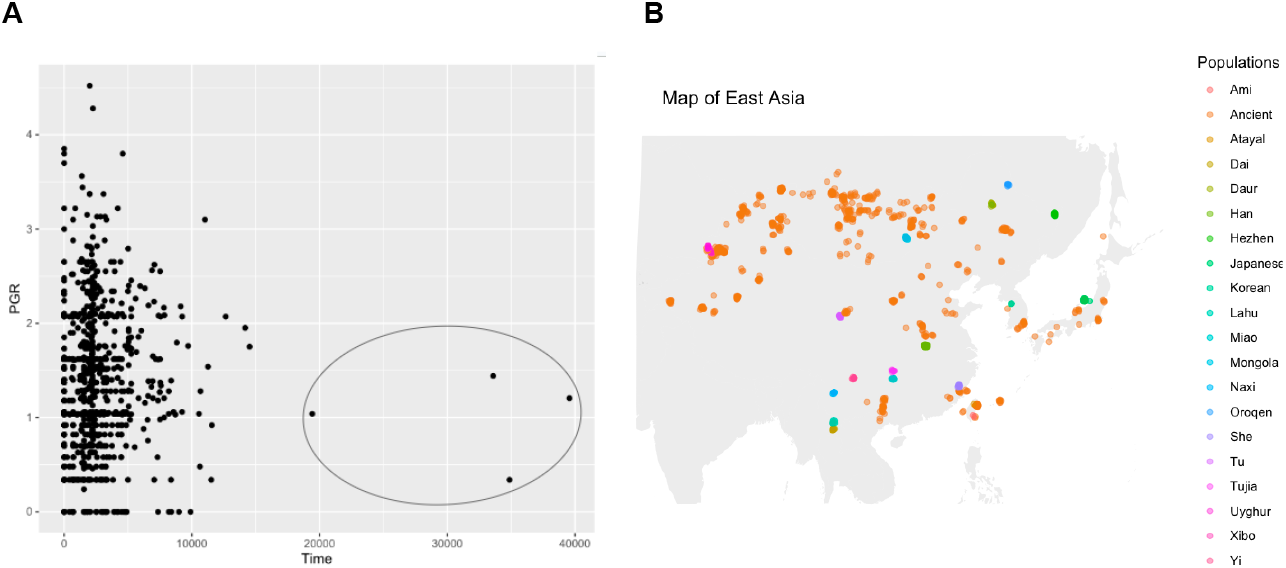
A) Change in PGR during the last 40.000 ya using 945 ancient and modern East asians. Each dot represents an ancient individual. All individuals older than the split among East Asians and Native Americans (inside the circle) have low PGR values, likely suggesting they had toned skin pigmentation. Individuals with a higher likelihood to develop lighter skin pigmentation dated for some thousands years after the split among East Asians and native Americans. B) Map of East Asia. All East Asian individuals, including modern and ancient, are plotted by the latitude and longitude indicating where they lived. Each dot represents an individual and each population is highlighted by a determined colour. Ancient individuals are coloured in orange.

## 4. Conclusion

Light skin tone is one of the most well studied human phenotypes, still there are many questions to be answered. On the other hand, its polygenic nature makes it difficult to study it using single variants. It has been suggested that it evolved in parallel in some populations like Europeans and East Asians. But, most studies came only from European populations (both using modern and ancient genomes), making other human populations to be underrepresented. Additionally, there is some controversy about when this phenotype evolved in East Asians, with some studies pointing to it being shared among East Asians and Native Americans and then being selected in the common ancestor of both populations. Whereas, others suggest a more recent origin. Although only some variants have been associated with light skin in East Asians, during the last years several articles have been published with ancient genomes encompassing the last 40,000 years. This allows us to study the evolution of this phenotype in East Asia through time. Here, I applied a Polygenic risk score (PGR) with light skin variants found in East Asians to shed light about when it may have evolved. Although the ancestors of both populations may have evolved some light skin tone, the results here point toward the phenotype strongly changed after the split among Native American and East Asians. However, only few variants associated with light skin have been found due to clear underrepresentation of East Asians genetic studies. These results have to be taken carefully, despite the significant difference in the PGR values among the different populations, the fluctuation inside each population is high and the variance among populations is still fine. This may be due to the fact that several populations show high interindividual variation, but also to the fact that only 7 SNPs are available to be used. However, I hope this study sets light on how to study complex phenotypes sub represented populations such as East Asians. I highly encourage future studies to focus on non representatives populations rather than studying only Europeans. For example, increasing the number of SNPs related to light skin in East Asians populations will allow us to conduct high accurate analysis to understand differences among populations, geographical variables and the evolution of skin tone.

## Acknowledgements

I would really like to thank Chen Liebson for her advice and her kindly support during the elaboration of the early manuscript.

## References

1. Dan, J. & Mathieson, I. 2021. The evolution of skin pigmentation-associated variation in West Eurasia. Proc. Natl. Acad. Sci. U.S.A. 118(1): e2009227118.

2. Jablonski, N.G. & Chaplin, G. 2010. Human skin pigmentation as an adaptation to UV radiation. PNAS. 117: 8962–8968.

3. Deng, L. & Xu, S. 2018. Adaptation of human skin color in various populations. Hereditas. 155:1. 10.1186/s41065-017-0036-2.

4. Neale Lab. UK Biobank GWAS. http://www.nealelab.is/uk-biobank.

5. Meyer, M., et al. 2012. A High-Coverage Genome Sequence from an Archaic Denisovan Individual. Science. 338: 6104.

6. Dalton, R. 2010. European and Asian genomes have traces of Neanderthal. Nature 10.1038/news.2010.225

7. Mathieson, I. et al. 2015. Genome-wide patterns of selection in 230 ancient Eurasians. Nature. 528: 499–503.

8. Norton, H.L., Kittles, R.A., Parra, E., McKeigue, P., Mao, X., Cheng, K., Canfield, V.A., Bradley, D.G., McEvoy, B. & Shriver, M.D. 2007. Genetic evidence for the convergent evolution of light skin in Europeans and East Asians. Mol. Biol. Evol. 24(3):710–22. doi: 10.1093/molbev/msl203.

9. Edwards, M., Bigham, A., Jonze, T., Li, S., Gozdzik, A., Ross, K., Jin, L. & Parra, E.J. 2010. Association of the OCA2 Polymorphism His615Arg with Melanin Content in East Asian Populations: Further Evidence of Convergent Evolution of Skin Pigmentation. PLoS Genet 6(3): e1000867. 10.1371/journal.pgen.1000867

10. Adhikari, K., Mendoza-Revilla, J., Sohail, A., Fuentes-Guajardo, M., Lampert, J., Chacón-Duque, J.C. et al. 2019. A GWAS in Latin Americans highlights the convergent evolution of lighter skin pigmentation in Eurasia. Nature. 10: 358.

11. Norton, H.L., Kittles, R.A., Parra, E., McKeigue, P., Mao, X. et al. 2007. Genetic Evidence for the Convergent Evolution of Light Skin in Europeans and East Asians, Molecular Biology and Evolution. 24 (3): 710–722, 10.1093/molbev/msl203

12. Seo, J.Y., You, S.W., Shin, J-G., Kim, Y., Park, S.G., Won, H.H. & Kang, N.G. 2022. GWAS Identifies Multiple Genetic Loci for Skin Color in Korean Women. Journal of Investigative Dermatology. 142 (2):1077–1084. 10.1016/j.jid.2021.08.440

13. Eaton, K., Edwards, M., Krithika, S., Cook, G., Norton, H. & Parra, E.J. 2015. Association study confirms the role of two OCA2 polymorphisms in normal skin pigmentation variation in East Asian populations. Am. J. Hum. Biol. 27 (4):520–525. 10.1002/ajhb.22678

14. Ferrando-Bernal, M. Ancient DNA suggests anaemia and low bone mineral density as the cause for porotic hyperostosis in ancient individuals. Sci Rep 13, 6968 (2023). 10.1038/s41598-023-33405-7

15. Igo, R. P. Jr., Kinzy, T. G. & Bailey, J. N. C. Genetic risk scores. Curr. Protoc. Hum. Genet. 104(1), e95 (2019).

16. Marciniak, S. et al. An integrative skeletal and paleogenomic analysis of stature variation suggests relatively reduced health for early European farmers. PNAS 119(15), e2106743119 (2022).

17. Cox, S. L. et al. Predicting skeletal stature using ancient DNA. Am. J. Biol. Anthropol. 177(1), 162–174 (2022).

18. Esteller-Cucala, P. et al. Genomic analysis of the natural history of attention-deficit/hyperactivity disorder using Neanderthal and ancient Homo sapiens samples. Sci. Rep. 10, 8622 (2020).

19. Purcell, S. et al. PLINK: A tool set for whole-genome association and population-based linkage analyses. Am. J. Hum. Genet. 81(3), 559–575 (2007).

20. RStudio Team. RStudio: Integrated Development for R http://www.rstudio.com/. x(RStudio, PBC, 2020).

21. Wickham, H. 2016. ggplot2: Elegant Graphics for Data Analysis. Springer-Verlag New York. ISBN 978-3-319-24277-4, https://ggplot2.tidyverse.org.

22. Tian, C., Kosoy, R., Lee, A., Ransom, M., Belmont, J.W., et al. 2008. Analysis of East Asia Genetic Substructure Using Genome-Wide SNP Arrays. PLoS ONE 3(12): e3862. 10.1371/journal.pone.0003862

23. Mehdi, J. et al. 2020. Investigating the Current State of Solar Energy Use in Countries with Strong Radiation Potential in Asia Using GIS Software, A Review Mehdi Jahangiria. Journal of Solar Energy Research Vol 5(3):477–497.

24. Willerslev, E. & Meltzer, D.J. 2021. Peopling of the Americas as inferred from ancient genomics. Nature. 594: 356–364.

25. Raghavan, M., Steinrücken, M., Harris, K., Schiffels, S., Rasmussen, S., et al. 2015. Genomic evidence for the Pleistocene and recent population history of Native Americans. Science. 349 (6250): DOI: 10.1126/science.aab3884

